# Litter characteristics and helping context during early life shape the responsiveness of the stress axis in a wild cooperative breeder

**DOI:** 10.1101/2021.06.11.448015

**Authors:** Aurélie Cohas, Coraline Bichet, Rébecca Garcia, Sylvia Pardonnet, Sophie Lardy, Benjamin Rey

**Author notes:** corresponding author: Benjamin Rey.

## Abstract

Stress responses have evolved to quickly and appropriately deal with environmental stressors in order to secure or restore homeostasis. Since the regulation of stress hormones plays a key adaptive role, the regulatory processes controlling stress hormones levels may be under high selective pressure. The social environment during early life (parents and litter characteristics) strongly affects ontogeny of the hypothalamic-pituitary-adrenal (HPA) axis. In cooperative breeders, offspring are also confronted with helpers but whether and how variation in the helping context can affect HPA axis responsiveness of offspring remains unanswered. Combining dexamethasone suppression and adrenocorticotropic hormone stimulation tests, we investigated the link between the social environment and the characteristics of the HPA axis at the early stages of life in wild Alpine marmots. We show that when raised in the presence of helpers, marmot pups exhibit a greater capacity not only to mount, but also to turn off a stress response. The capacity to mount a stress response was also higher as the pups were raised in large litters. Determining impacts of such social modulation of the HPA axis functioning on individual fitness would make an important contribution to our understanding of the evolution of cooperative breeding.

## 1. Introduction

Organisms constantly adjust to predictable environmental variations such as daily and seasonal changes while needing to respond quickly and adequately to unpredictable treats (Wingfield 2003, 2008). To do so, stress responses have evolved to deal rapidly and proportionately with environmental stressors in order to maintain or restore a homeostatic state (Monaghan and Spencer 2014). The stress response consists of a complex set of physiological changes through the activation of neuronal and endocrine pathways (Sapolsky et al. 2000; Wingfield 2003) where the hypothalamic-pituitary-adrenal (HPA) axis is the main orchestrator by regulating the secretion and release of glucocorticoids (de Kloet et al. 2008). At baseline levels, circadian fluctuations of glucocorticoids (corticosterone and cortisol) regulate physiological functions and maintain energy homeostasis (McEwen and Wingfield 2003). When exposed to physiological or psychological stressors, activation of the HPA axis stimulates the secretion and release of glucocorticoids within minutes, mobilizing energy and preparing individuals to fight or flight (Wingfield et al. 1998). While a transient increase in glucocorticoids leads to short-term and immediate benefits, particularly in terms of survival (Wingfield 2008), chronically elevated glucocorticoid levels carries physiological costs (Romero et al. 2009). Under high glucocorticoids levels, energy is derived to an emergency state, where physiological and behavioural adjustments are all directed towards immediate survival, at the expense of other energy demanding functions such as growth, reproduction or body maintenance (Romero 2004). Extensive literature, mostly on medical studies in humans and experimental approaches on laboratory rodents, agree that maintaining high levels of glucocorticoids cause adverse effects on many vital physiological functions while predisposing to multiple metabolic and immune disorders and accelerating ageing (for example, see Gassen et al. 2017; Nicolaides et al. 2015). In the wild, chronic elevation of glucocorticoids has been shown to have consequences on individual fitness with measurable cascading effect at the population level (eg. Boonstra et al. 1998; Dulude-de Broin et al. 2020). Hence, regulatory mechanisms of the stress response which control both the ability to mount a stress response (the initial response phase to challenge) and the mechanisms contributing to restore baseline glucocorticoid levels (the recovery phase), should play a key adaptive role (Crespi et al. 2013; Taff and Vitousek 2016) and thus under high selective pressure (MacDougall-Shackleton et al. 2013; Bonier and Martin 2016).

The stress response is driven by the activation of the HPA axis, which results in the hypothalamus releases of corticotropin-releasing hormone (CRH), a neuropeptide that stimulate the secretion of adrenocorticotropic hormone (ACTH) by the anterior pituitary gland. ACTH is then transported through bloodstream and stimulates the systemic release of glucocorticoids by the adrenal cortex (Figure 1a). The adrenocortical reactivity to ACTH is a major determinant of glucocorticoid elevation in blood and therefore controls the rate of the increase in energy demand and changes in resources allocation (i.e. emergency life-history stage) during the stress challenge (Wingfield et al. 1998). Conversely, the primary inhibitory control of HPA axis involves glucocorticoid-mediated negative feedback on the release of CRH and ACTH, so that the glucocorticoid levels return to baseline once the threat is dealt with. The reactivity of the adrenal to ACTH, as well as the effectiveness of negative glucocorticoid feedback, are plastic physiological traits whose expression aims to optimize the intensity and duration of stress responses in a given context (Angelier and Wingfield 2013). Although evidences that the reactivity of HPA traduces into to fitness benefits remains mixed (see Breuner et al. 2008), it is likely to be under high selective pressure (Bonier and Martin 2016).

**Figure 1.**
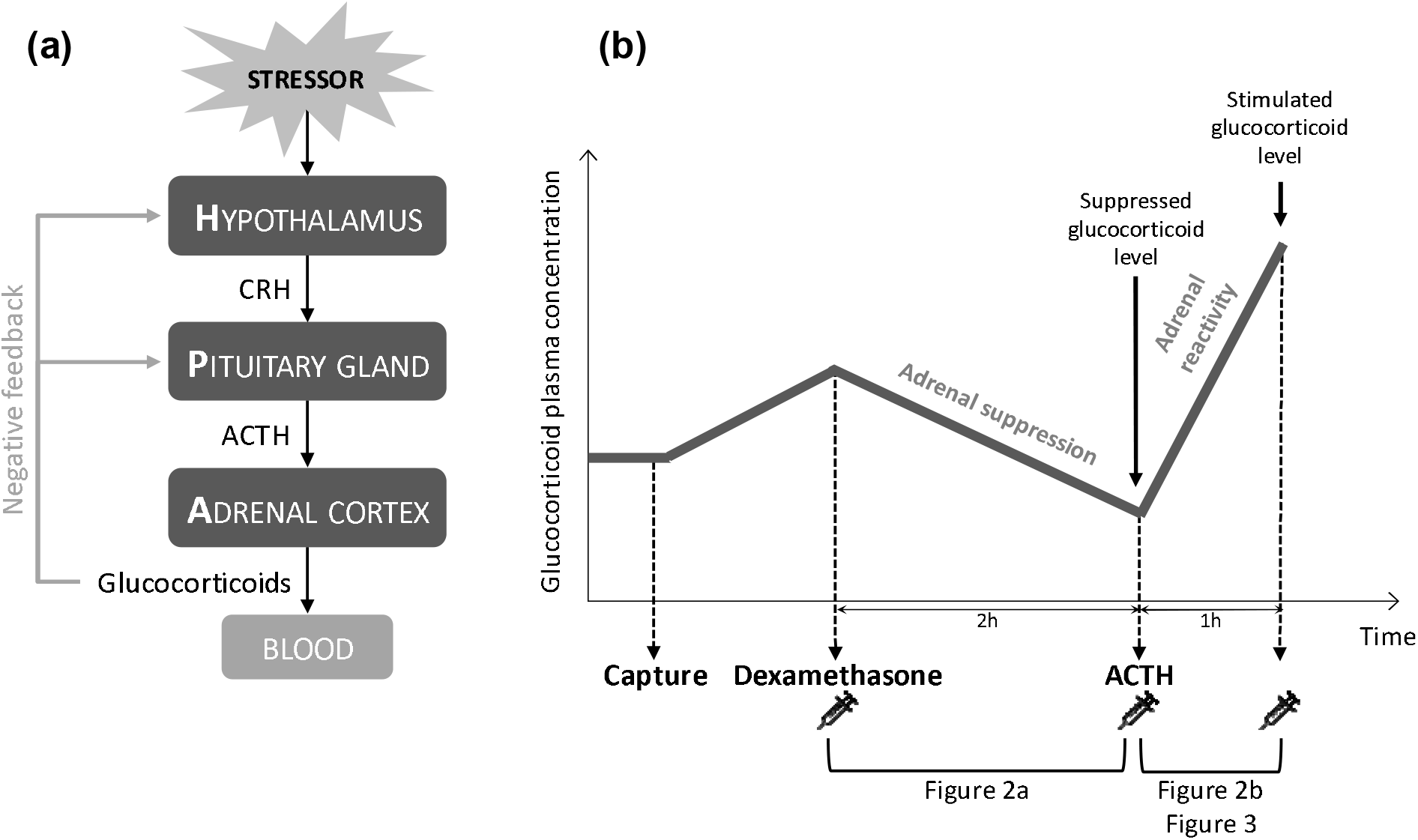
(a) Schematic representation of the stress response through the hypothalamic-pituitary-adrenal (HPA) axis. The stressor may be environmental, physical or psychological. It causes secretion of corticotropin-releasing hormone (CRH) by the hypothalamus. Under CRH stimulation, the pituitary gland releases adrenocorticotropic hormone (ACTH) into the bloodstream that in turn stimulates glucocorticoid secretion by the adrenal gland. Glucocorticoids retroact at the hypothalamus and pituitary gland levels inhibiting CRH and ACTH release. Following HPA axis stimulation, if the negative feedback mechanism works efficiently, plasma stress hormones rapidly returns to baseline level. (b) Illustration of the experimental protocol. Plasma glucocorticoid concentration rises following capture. Injection of dexamethasone leads to the suppression of the glucocorticoid secretion by the adrenal gland. The rate of dexamethasone-induced decrease of plasma glucocorticoid probes the efficiency of the negative feedback and the ability of the HPA axis to return to baseline level. The injection of ACTH, on the other hand, triggers a strong transient increase in plasma glucocorticoid concentration, an effect that reveals the reactivity of the adrenal gland.

Both the abiotic and biotic components of the environment can affect individual stress response (Creel et al. 2013; Grace and Anderson 2018). Experimental approach shows that when experienced early in life, environmental conditions strongly affect the ontogeny of the HPA axis, resulting in long-term effects on many physiological and neural functions (McMillen and Robinson 2005), and contributing greatly to a phenomenon referred as “developmental programming” (Seckl 2004). For instance, in rats, offspring that received extensive maternal care in early life showed reduced adrenal reactivity to acute stress, but improved efficacy of glucocorticoids negative-feedback in adulthood (Liu et al. 1997).

The social environment early in life is known to have an impact on glucocorticoids levels although most of our knowledge in this field comes from experimental approaches. For example, social deprivation (social isolation: (Malkesman et al. 2006); deprivation of mother and siblings: (Avishai-Eliner et al. 1995)) or social crowding experienced early in life (Bugajski et al. 1993) have been shown to cause chronic stress in laboratory rodents. Under natural conditions, investigations on the effects of the social environment in the early life have showed that litter size and composition affected circulating glucocorticoid of rat and rabbit pups (Hudson et al. 2011). However, in social species, offspring interact not only with their parents or siblings, but also with other members of their social groups. In particular, in cooperative breeders, animals live in kin-based family groups where it is not only the parents who care for the young (Brown 1987), but also non-breeding individuals called helpers (Keller and Reeve 1994). The presence of helpers early in life has major short- and long-term effects on offspring’s phenotypes and fitness (*e.g.* Berger et al 2018; Hammers et al 2019), but the underlying physiological mechanisms are not yet understood. In this study, we tested the hypothesis that the presence of helpers could modulate the HPA responsiveness of the offspring they care for.

In order to investigate whether the social environment (litter characteristics and helping context) experienced early in life shapes the individual HPA axis responsiveness, we implemented hormonal challenge protocols in the wild Alpine marmot (*Marmota marmota*). Alpine marmots are territorial cooperatively breeding ground-dwelling squirrels that live in family groups of 2 to 16 individuals composed of a dominant pair and sexually mature and immature subordinates (Allainé 2000). Only the dominant pair reproduces once a year. Subordinates of both sexes delay dispersal and forgo their own reproduction beyond sexual maturity (Cohas et al. 2006). However, only subordinate males are considered as helpers, notably because of their crucial role in thermoregulation which drastically improves the survival of juveniles during hibernation (Arnold 1988; Allainé and Theuriau 2004) but also because of their role in the thermoregulation of the pups at night and in the construction of nests. Mating takes place in the second half of April, gestation lasts 35 days, the females then give birth to altricial pups and nurse them for 40 days. The pups remain in their natal underground burrows (i.e. in a buffered environment offering protection from climatic fluctuations and predators) until weaning, from the last week of June to mid-July, at about 40 days of age. During the whole nursing period, the impacts of group characteristics on these naïve individuals are thus expected to be prominent. We previously showed (1) that litter sex composition and size affect male juvenile Alpine marmot survival and probability to become dominant for both sexes (Dupont et al. 2015), and (2) that the number of helpers during early life has long-term consequences such as increased longevity of females (Berger et al. 2015) or alteration of actuarial senescence pattern in both sexes (Berger et al. 2018). However, whether litter size, sex-ratio and helpers can impact pups physiological traits remains to be investigated. From these previous studies, we predict that the pups benefiting from helpers early in life should present increased HPA axis responsiveness with enhanced capacity to rapidly mount and to turn off a stress response. We also expect the litter size and sex-ratio to affect the HPA axis responsiveness.

## 2. Materials and Methods

### (a) Field methods

The study took place in La Grande Sassière Nature Reserve (French Alps, 45°29’N, 65°90’E) where a population of wild Alpine marmots (*Marmota marmota*) is extensively monitored since 1990. As part of a long-term capture-mark-recapture protocol, marmots are captured annually from mid-April to mid-July using two-door live-capture traps (see Cohas et al. 2006). Once a year, captured individuals are tranquilized by an intramuscular injection of tiletamine and zolazepam (Zolétil 100, 0.1 ml.kg^−1^, Virbac, France), sexed, aged, and their social status (dominant or subordinate) is determined by examining sexual characteristics (scrotum for males and teats for females). Social status is further confirmed by behavioural observations. Each animal is individually marked with a microchip (Trovan, Germany) inserted subcutaneously and a numbered ear-tag. Dominant individuals also received a coloured plastic ear-tag.

Fieldwork work was undertaken after the issuance of permit number AP n82010/121 by the Prefecture of Savoie. All the procedures were approved by the ethical committee of the University Claude Bernard Lyon 1 (n8BH2012-92 V1, 2017012500169084 v1).

### (b) Characterization of the social environment

We observed each family group on average 1h per day for a minimum of 30h per year with sessions being randomly distributed during the activity period. Marmots were observed from a distance of 80-200 m using 10×50 binoculars and 20×60 monocular telescopes. Ear-tags allowed us to distinguish the sex and social status of individuals and their size allowed us to categorize them as yearlings, 2-year-olds or adults. Combining these observations with mark-recapture data, we defined the number of helpers as the number of male subordinates aged 1 year and older present in a family group from the end of mating to the emergence of the pups. The number of helpers (median = 1.50, range = 0-4) was not manipulated, but we benefited from the natural intergroup variability in the number of helpers in this population. Among the 45 pups studied, 12 were raised in absence of helpers in their family group.

From additional daily observations conducted during the whole activity period, we recorded the date of the pups’ first emergence from the natal burrow and the litter size. All pups of a given litter emerge together. We defined the litter size (median = 4.00, range = 2-6) as the total number of pups emerging from the same natal burrow. Combining observations with the capture of the pups, we characterized the composition of the litter by the litter sex-ratio (median = 0.75, range = 0.25-1), the litter sex-ratio being measured as the number of males in a litter divided by the litter size. Sibship was further confirmed by parentage analyses (Cohas et al. 2006).

### (c) Corticosterone assay

In vertebrates, two adrenal glucocorticoids are secreted: cortisol and corticosterone. While these two hormones are believed to play a role in the stress response of vertebrates, one is generally predominant and the cortisol to corticosterone ratio is varies considerably depending on the species (Koren et al. 2012). In rodents, it is generally accepted that the dominant glucocorticoid is corticosterone (Cockrem, 2013), and this hormone or its derivatives has been preferred to cortisol in literature on marmot species (eg. Monclus et al. 2011; Petelle et al. 2017; Blumstein et al. 2018; Price et al. 2018; Pinho et al. 2019). However, there is no consensus on this point since other authors have shown higher cortisol than corticosterone concentrations in yellow-bellied marmot plasma (Kastner et al. 1977). Here, we measured total (bound and unbound) corticosterone in plasma using commercial enzyme immuno-assay kits (Cayman Chemicals Corticosterone EIA kits n°501320). The assays were carried out in microplates coated with mouse anti-rabbit IgG and relied on the competition between corticosterone and a corticosterone-acetylcholinesterase for a limited amount of corticosterone antiserum. Corticosterone-acetylcholinesterase complexed with the IgG colours the Ellman’s reagent in yellow, the intensity of which is measured spectrophotmetrically at 412nm (Xenius, Safas Instrument) and is inversely proportional to corticosterone in the sample. This assay has been validated for rodents (see manufacturer’s instructions), nevertheless, we checked the specificity of the corticosterone assay by verifying that the hormone levels measured in a series of plasma diluted in assay buffer (from 1:5 to 1:20) were parallel to the dilution series of the hormonal standards. Final assays were performed in duplicate with a volume of 50μL of plasma diluted 1:10. The mean intra-assay coefficient of variation that was calculated for each sample was 6.8%. The corticosterone concentrations were calculated against a standard curve and expressed as pg.mL^−1^.

### (d) Characterization of the stress axis

In 2015, 45 pups from 13 family groups were captured by hand between their first and third day of emergence from their natal burrow (0 to 2 days after emergence). At emergence, pups are weaned and beyond the stress hyporesponsive period, which is known to be in the first half of the period between birth and weaning in rodents (Levine 1994). Immediately upon capture, the pups were transferred to a nearby field laboratory in a burlap bag, before being tranquilized and tagged as described above. To limit potential confounding effects of environmental variability, the experiments were all conducted the same year in family groups occupying valley and south facing territories. Given the size of the overall study site (∼0.3 km^2^) and the low range of altitude (∼50m), all studied territories were expected to face very similar climatic conditions.

To assess measures of pup’s stress axis responsiveness and resulting corticosterone levels we recorded complementary measures. Firstly, we applied a dexamethasone suppression test (Boonstra et al., 1998). Dexamethasone is a synthetic steroid molecule that exerts a negative feedback on the hypothalamus and the anterior pituitary gland inhibiting the secretion of CRH and ACTH thus suppressing glucocorticoids release by the adrenal cortex (Fig 1a). Therefore, measuring the rate of plasma corticosterone disappearance following dexamethasone administration 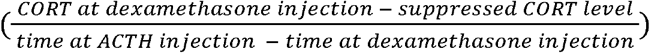, hereafter referred as “adrenal suppression by dexamethasone”) provides an effective measure of the efficacy of the negative feedback regulation. In other words, it measures the ability of the HPA to recover from initial stimulation and to return to baseline level. A low efficiency of negative feedback indicates a reduced ability to shut-off the release of glucocorticoids, exposing individuals to the adverse effects of prolonged stress. Such an effect has been observed repeatedly in wild and captive vertebrate populations subjected to persistent stress (see Dickens and Romero 2013 for review). Corticosterone concentration measured two hours post-dexamethasone injection (hereafter referred as “suppressed corticosterone level”) assumes effective inhibition of endogenous CRH and ACTH secretion. Secondly, we performed an ACTH stimulation test (Boonstra et al., 1998). Following exogenous ACTH administration, the appearance of the corticosterone in the blood 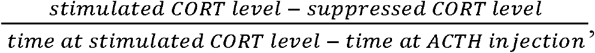, hereafter referred as “adrenal reactivity to ACTH”) measures the efficiency of the adrenal cortex to produce and release stress hormones (Fig 1a). In turn, this test informs directly on the reactiveness of the adrenal gland and on its ability to initiate glucocorticoids-dependant adaptive stress responses (Boonstra et al. 1998), a faster glucocorticoid release after ACTH stimulation indicating a greater physiological capacity to orchestrate an effective response to a perceived stressor. The corticosterone concentration measured one hour after ACTH injection (hereafter referred to as “stimulated corticosterone level”) reflects the secretory activity of the adrenal gland during that time under ACTH stimulation.

Each pup thus underwent three successive blood sampling (Fig 1b) of 200μl consisting in a small puncture of the great saphenous vein with a 0.5 × 16mm sterile needle (Terumo, Europe). Blood was collected in heparinized capillary tubes (Microvette, Sarstedt, Germany). Immediately upon the first blood sampling, pups received an intramuscular injection of dexamethasone sodium phosphate (D0720000, Sigma-Aldrich, France, 0.4 mg.kg^−1^). A second blood sample was taken two hours (mean±SD = 1:59±0:01, range = 1:55-2:03) after dexamethasone administration and was immediately followed by an intramuscular injection of ACTH (Sigma-Aldrich, France, 4.0 IU.kg^−1^). A third blood sample was collected one hour (mean±SD = 0:59±0:01, range = 0:51-1:08) after ACTH injection. The times between dexamethasone or ACTH injections and blood sampling were determined from a preliminary trial on 5 individuals for whom multiple blood samples were taken at different time interval after injections (30 min, 1h, 2h, 3h). For all individuals, two hours were required to obtain a marked decrease in plasma corticosterone. For 3 of the 5 individuals tested, peak corticosterone level was already reached and corticosterone concentration decreased two hours after ACTH injections; we thus set the blood sampling at 1 hour after the ACTH injection in or protocol. All blood samples were centrifuged for 5 min at 3000×g and the plasma stored in the field at −20°C. At the end of the procedure, pups were released at their natal burrow. They were observed on a daily basis for 3 days; all individuals were reintegrated to their family group without any adverse effects being observed.

### (e) Statistical analyses

To test whether social environment affects corticosterone levels and HPA axis responsiveness of marmot pups, the suppressed and the stimulated corticosterone levels as well as the adrenal suppression by dexamethasone and the reactivity of the adrenal gland to ACTH were entered as dependent variables respectively in two generalized linear mixed models with a logarithm link and a variance given by a gamma distribution and in two linear mixed models (LMM). To test for an effect of litter characteristics, the litter size and sex-ratio were entered as explanatory variables. To test for an effect of the helping context, the number of helpers was entered as an explanatory variable. Additionally, the sex and the age (in day since emergence from the natal burrow) of the pup were entered as potential confounding explanatory variables. To control both for pseudo-replication arising from measures done for several pups of the same litters sharing a common environment (*e.g.* genetic background, maternal effects, territory quality), we entered a variable “litter” as a random intercept. Agreeing with our hypotheses, only additive effects of all the fixed explanatory variables were considered in our models. Although inspection of the residuals from models including only previous effects did not show any evidence of potential interaction between the considered explanatory variables, to assure the robustness of the results obtained from these previous models, we further constructed models with all 2-way interactions. Interactions were removed if not significant following a backward procedure. Both approaches gave the exact same results.

Statistical analyses were performed with R 3.3.1 (R Core Team, 2014). The function ‘‘lmer’’ in the package ‘‘lme4’’ (Bates et al. 2015) was used to fit GLMMs (Venables and Ripley 2002) and the package “lmerTest” (Kuznetsova et al. 2016) used to calculate parameter specific *p*-values based on Satterthwaite’s approximations. We set the level of significance to α = 0.05 and parameter estimates are given as mean ± SE.

## 3. Results

The average suppressed and stimulated corticosterone levels were 1837 ± 877(SD) and 4080 ± 1703(SD) pg.ml^−1^ respectively (See Supplementary Information). The average adrenal suppression by dexamethasone and adrenal gland reactivity to ACTH were −9.42 ± 10.70 (SD) pg.ml^−1^.min^−1^ and 37.37 ± 19.81(SD) pg.ml^−1^.min^−1^ respectively (see supplementary material). Adrenal reactivity to ACTH, adrenal suppression and the suppressed and stimulated corticosterone levels showed some degree of correlation (Table 1).

**Table 1.** Pearson coefficients of correlation between the different corticosterone levels measured and both adrenal reactivity to ACTH and adrenal suppression by dexamethasone. Significant correlations are in bold (p<0.05).

|  | Suppressed<br>corticosterone level | Stimulated<br>corticosterone level | Adrenal suppression<br>by dexamethasone | Adrenal reactivity to<br>ACTH |
| --- | --- | --- | --- | --- |
| Corticosterone at<br>dexamethasone<br>injection | **0.65**<br>**95% CI[ 0.43;0.79]**<br>**t = 5.54**<br>**N = 45**<br>**p < 0.01** | **0.68**<br>**95% CI[ 0.48;0.82]**<br>**t = 6.01**<br>**N = 43**<br>**p < 0.01** | **-0.85**<br>**95% CI[ -0.92;-0.74]**<br>**t = 10.67**<br>**N = 45**<br>**p < 0.01** | **0.51**<br>**95% CI[ 0.24;0.70]**<br>**t = 3.75**<br>**N = 43**<br>**p < 0.01** |
| Suppressed<br>corticosterone<br>level |  | **0.76**<br>**95% CI[ 0.59;0.87]**<br>**t = 7.55**<br>**N = 43**<br>**p < 0.01** | -0.15<br>95% CI[-0.43;0.16]<br>t = 1.00<br>N = 45<br>p = 0.33 | **0.38**<br>**95% CI[0.08; 0.62]**<br>**t = 2.63**<br>**N = 43**<br>**p = 0.01** |
| Stimulated<br>corticosterone<br>level |  |  | **-0.38**<br>**95% CI[-0.62;-0.09]**<br>**t = 2.67**<br>**N = 43**<br>**p = 0.01** | **0.89**<br>**95% CI[0.79;0.94]**<br>**t = 12.29**<br>**N = 43**<br>**p < 0.01** |
| Adrenal<br>suppression by<br>dexamethasone |  |  |  | **-0.41**<br>**95% CI[-0.63;-0.11]**<br>**t = 2.84**<br>**N = 43**<br>**p < 0.01** |

The suppressed and the stimulated corticosterone levels of the pups were not affected by the number of helpers in their family groups or the litter sex-ratio (Table 2). The stimulated corticosterone level was significantly affected by the litter size but not the suppressed corticosterone level (Table 2).

**Table 2.** Effect of the litter size and composition as well as of the number of helpers on HPA responsiveness and plasma corticosterone levels of Alpine marmot pups subjected to dexamethasone suppression and ACTH stimulation tests. Significant effects are in bold (p<0.05).

| Fixed effects | Adrenal suppression by dexamethasone<br>N = 45 |  |  | Suppressed corticosterone level<br>N = 45 |  |  | Adrenal reactivity to ACTH<br>N = 43 |  |  | Stimulated corticosterone level<br>N = 43 |  |  |
| --- | --- | --- | --- | --- | --- | --- | --- | --- | --- | --- | --- | --- |
|  | Estimate ± SE | t value | p-value | Estimate ± SE | t value | p-value | Estimate ± SE | t value | p-value | Estimate ± SE | t value | p-value |
| Intercept | 8.01 ± 10.92 | 0.73 | 0.48 | 1329 ± 1055 | 1.26 | 0.23 | -10.05 ± 13.93 | 0.72 | 0.47 | -370 ± 1335 | 0.23 | 0.78 |
| Age | 0.44 ± 2.52 | 0.18 | 0.87 | 260 ± 217 | 1.20 | 0.24 | -6.27 ± 4.20 | 1.49 | 0.24 | -302 ± 402 | 0.75 | 0.46 |
| Sex (male) | 0.74 ± 2.97 | 0.25 | 0.81 | -95 ± 247 | 0.39 | 0.70 | -0.96 ± 5.55 | 0.17 | 0.86 | -20 ± 532 | 0.04 | 0.97 |
| Sex-ratio | -7.97 ± 9.36 | 0.85 | 0.41 | 1206 ± 911 | 1.32 | 0.21 | 17.98 ± 12.48 | 1.44 | 0.16 | 1973 ± 1196 | 1.65 | 0.11 |
| Litter size | 1.94 ± 2.33 | 0.83 | 0.42 | -16 ± 224 | 0.07 | 0.84 | **6.65 ± 3.08** | **2.16** | **0.04** | **623 ± 296** | **2.11** | **0.04** |
| Number of helpers | **-2.98 ± 1.29** | **2.31** | **0.04** | 11 ± 126 | 0.09 | 0.93 | **5.50 ± 1.63** | **3.36** | **0.002** | 276 ± 157 | 1.76 | 0.09 |

Both the adrenal suppression by dexamethasone (negative feedback, Fig 2a) and the adrenal gland reactivity to ACTH (Fig 2b) increased with the number of helpers (Table 2). The litter sex-ratio had no effect on both components of the HPA responsiveness but the litter size positively affected the adrenal gland reactivity to ACTH in pups (Table 2, Fig 3).

**Figure 2.**
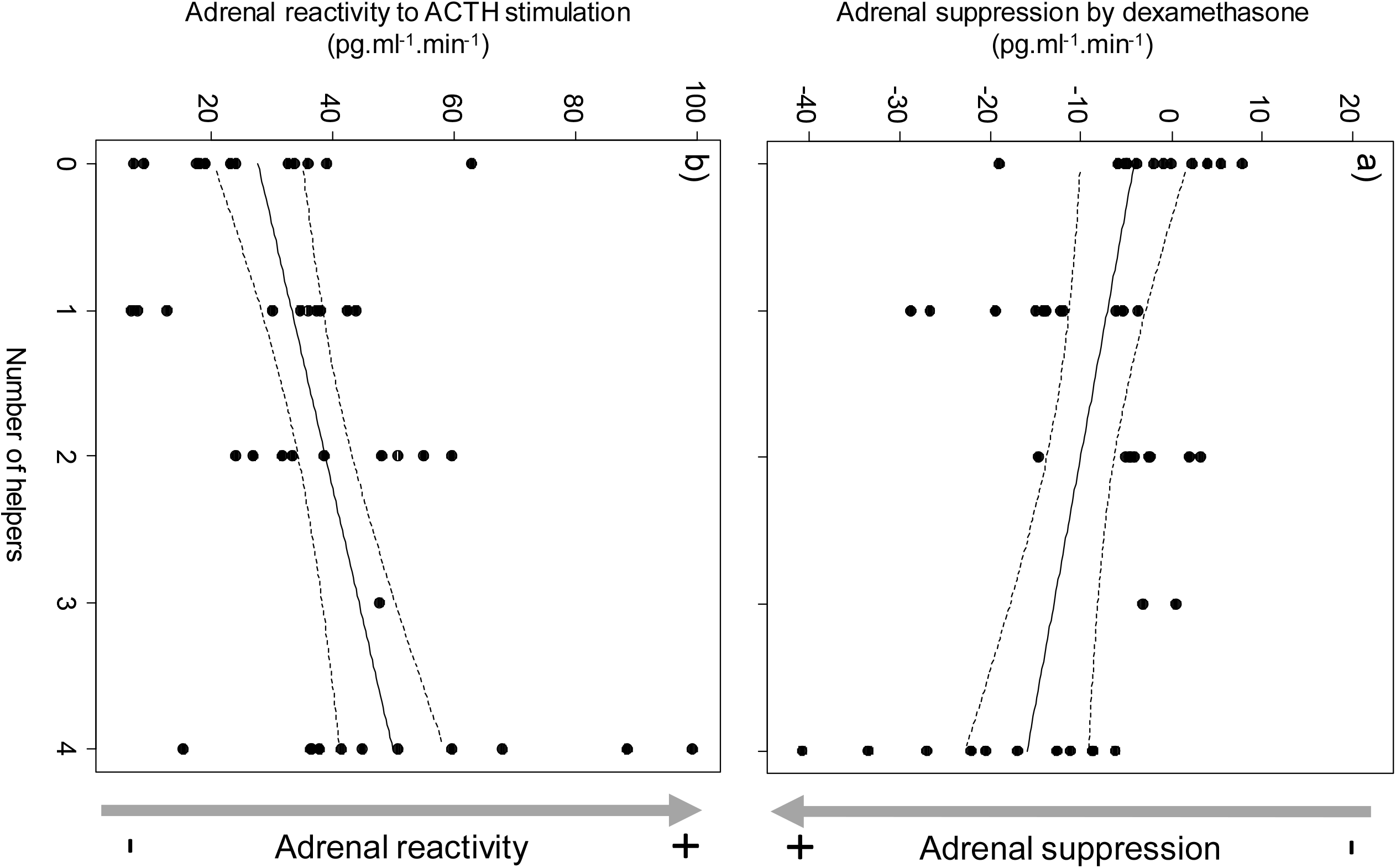
Effects of the number of helpers (a) on the adrenal suppression by dexamethasone (measured as plasma corticosterone disappearance following dexamethasone administration) and (b) on the adrenal reactivity to ACTH (measured as plasma corticosterone appearance following ACTH administration) of Alpine marmot pups. Black dots represent residuals (a) of the adrenal suppression by dexamethasone or of (b) adrenal reactivity to ACTH per number of helpers after controlling for other effects. Line represents the model predictions (black) and their associated standard errors (dotted).

**Figure 3.**
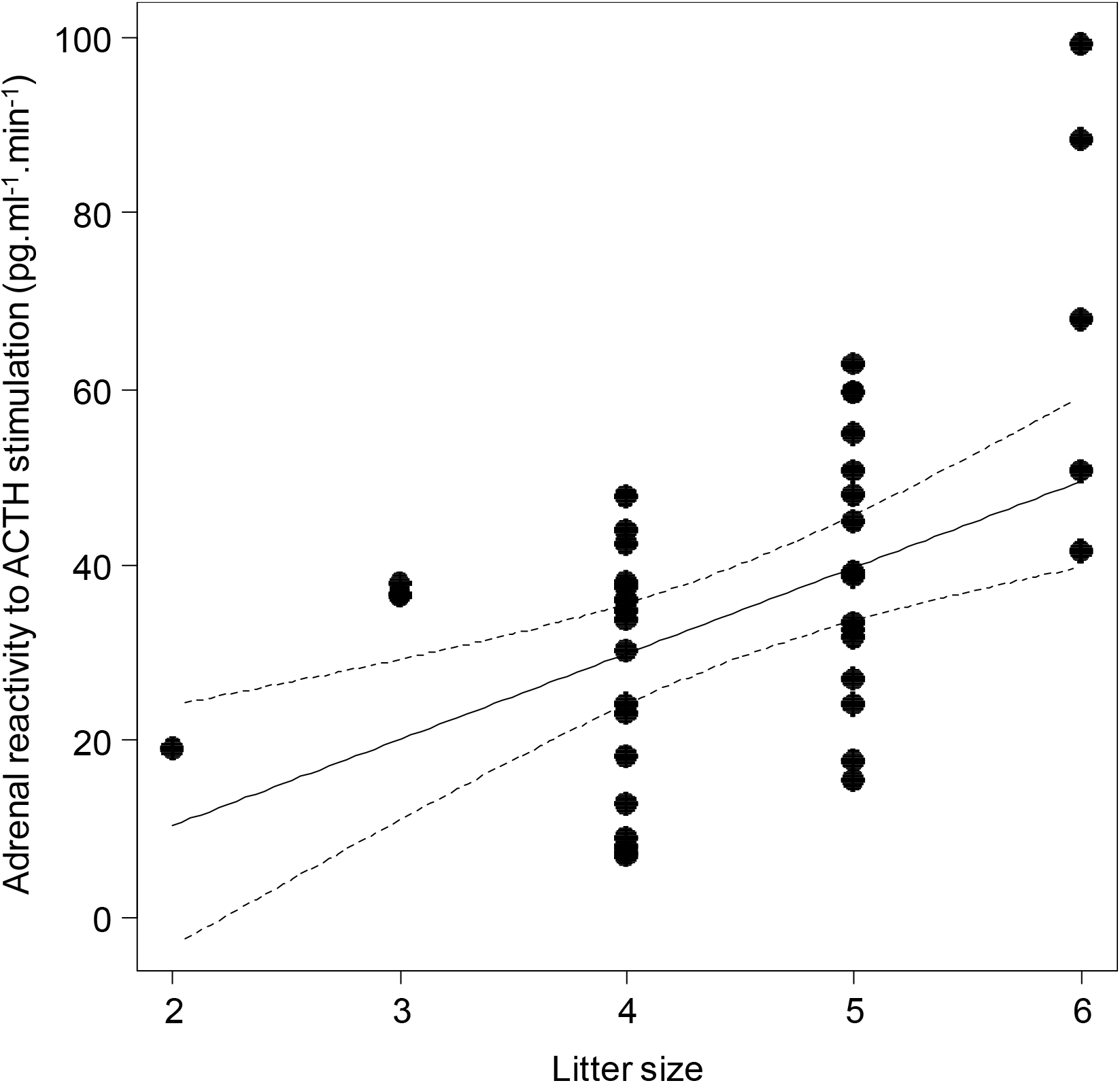
Effects of the litter size on the adrenal reactivity to ACTH of Alpine marmot pups. Black dots represent residual adrenal reactivity to ACTH after controlling for the effects of the number of helpers. Line represents the model predictions (black) and their associated standard errors (dotted).

## 4. Discussion

We showed that marmot pups raised in naturally contrasting social contexts display different sensitivity profiles of the HPA axis. An increase in litter size has led to a greater reactivity to mount a stress response and higher corticosterone level after stimulation by exogenous ACTH. While the helping context did not impact the corticosterone levels after stimulation by ACTH or inhibition by dexamethasone, the number of helpers significantly affected the pup’s HPA axis responsiveness: the presence of helpers triggered a higher reactivity of the adrenal gland to ACTH and a higher hypothalamo-pituitary sensitivity to inhibition by dexamethasone. These data indicate that, when raised in the presence of helpers, marmot pups thus exhibit a greater capacity not only to mount but also to turn off a stress response. Although the four measured physiological traits are not totally independent and showed some degree of correlations, they seemed to have been shaped by different evolutionary pressures linked to the social context.

We found that litter size but not litter sex-ratio had a significant impact on marmot pups’ adrenal reactivity to ACTH and stimulated corticosterone levels. Such effect is congruent with extensive literature showing variation in hormones levels linked to litter size. For instance, in small laboratory mammals (Fey and Trillmich 2008; Roedel et al. 2010), pups from large litters exhibit higher baseline glucocorticoids levels than pups from small litters due to increased competition during lactation, particularly in species where the number of offspring exceeds the number of teats. In contrast, the lack of sex-ratio effect is somehow surprising. Indeed, the sex composition of litters has been shown to influence the stress hormone levels of offspring (Benhaiem et al. 2013; Blanco et al. 2006). Both hormonal infusion during pregnancy and competition between siblings had been advanced to modulate the stress hormone levels (Blanco et al. 2006; Hudson et al. 2011; Benhaiem et al. 2013). Regarding the lack of effect of litter size and sex-ratio on adrenal suppression and suppressed corticosterone levels, comparison with the literature is not possible as no study to date have investigated the impact of litter characteristics on adrenal negative feedback.

As predicted, the HPA characteristics depended on the number of helpers present in a family group. This could arise through direct effects on pups. In Alpine marmots, as in other cooperative breeders, helpers provide important social support to offspring through babysitting (Clutton-Brock et al. 2000), food provisioning (Brotherton et al. 2001; Raihani and Ridley 2008), grouping or social thermoregulation (Arnold 1988). Hence, the beneficial effect of helping can be expected to allow pups to invest more in somatic functions and to accelerate the ontogeny of physiological traits such as the HPA axis. If there is no evidence that helping influence the age at emergence from the natal burrow, helping behaviours clearly enhance offspring phenotypic quality, notably growth and body mass (Clutton-Brock et al. 2001; Hodge 2005). Despite nothing is known about an alteration of the HPA axis functioning linked to help received directly as pups in cooperatively breeding species, the enhanced adrenal reactivity and feedback efficiency we observed in Alpine marmot pups could arise as benefits of help directed toward pups on HPA axis ontogeny and rate of maturation. Similarly, social support in humans (Uchino et al. 1996) and affiliative behaviours in other mammalian species (Hennessy et al. 2009; Tuchscherer et al. 2016) have been shown to affect HPA axis positively.v

Effect of helpers could also be indirect, for example by affecting the hormonal status of the mother. Indeed, maternal effects are a major determinant of offspring HPA axis profile and previous findings on vertebrates suggest that maternally-induced stress can cause significant variation in the responsiveness of an offspring’s HPA axis involving both pre- (Moisiadis and Matthews 2014) and postnatal developmental mechanisms (Liu et al. 1997; Macri and Wuerbel 2006). Helping context has been shown to modify both maternal condition and physiology leading to change in maternal allocation in birds (Emlen et al. 1986; Hatchwell 1999; Paquet et al. 2013; Dixit et al. 2017). In mammals, a reduced number of helpers raises hormone stress levels of the mothers (Cameron 2004). Effects of helpers in modifying maternal glucocorticoids levels could thus affect the functioning of the HPA axis in pups.

Finally, while the litter and the characteristics of the social group affected the reactivity of the pituitary-adrenal system of the pups, little or no effect was observed on the corticosterone levels after stimulation by ACTH or inhibition by dexamethasone. These results are in line with the previously stated idea that static measures of stress hormone levels do not always allow capturing information on overall endocrine functions. For instance, glucocorticoid value measured at the peak of secretion not only poses the problem of whether the peak has been properly identified, but is may be a poor indicator of the integrated stress response over time (Romero 2004).

Our approach also has limitations. Our protocol requires individuals to be tranquilized and constrained for several hours, which is hardly feasible for many wild species in their natural environment. The potential effect of the tranquilizing drug (Zoletil), which consisted in a combination of tiletamine (a dissociative anesthetic) and zolazepam (a benzodiazepine) should also be considered. Dissociative aesthetics have little or no adverse effect on the HPA axis functioning, however, but data on humans showed that benzodiazepines may compromise adrenocortical steroidogenesis in a dose-dependent manner (Besnier et al. 2016). Although the effect of zolazepam on HPA has yet to be demonstrated in wildlife, particularly at the dose used, and the doses were standardized per g of body weight to allow comparisons between individuals, it cannot be formally excluded that the tranquilization impacted our results. While our protocol allowed to evaluate the propensity of the adrenal gland to release of glucocorticoids into the bloodstream, there are other regulatory mechanisms that modulate the amplitude and the duration of the stress response. These include those affecting the pharmacokinetics of glucocorticoids (e.g. hepatic glucocorticoids metabolism, plasma clearance and excretion (McKay and Cidlowski 2003), and the concentration of binding proteins (Breuner and Orchinik 2002). Applying a holistic approach has become a necessary need to better understand the adaptive value of the responses to stress (Rey 2020). Hence, further investigations should focus on whether the social context impact other regulatory mechanisms such as concentration of glucocorticoids bindings proteins or the fraction of free hormone that is available for uptake by tissues. Further studies are also needed to clarify how modulation of the reactivity of the HPA axis by the social context translates into fitness and by which mechanisms (Breuner et al. 2008).

At this stage, we know that early social environment has major repercussions on an individual’s fitness (Beckerman et al. 2002; Lindström 1999). In Alpine marmot, help received during early life modulates female lifetime reproductive success (Berger et al. 2015) and both male and female actuarial senescence (Berger et al. 2018). Moreover, litter sex composition affects male juvenile survival and both male and female probabilities of reaching dominant status (Dupont et al. 2015). These very same social factors were the ones we found to modulate the reactivity of the HPA. Whether these factors, and more particularly helpers in cooperative breeders, could influence such long-lasting effects through an early programming of the HPA axis remain to be investigated and the following questions need to be answered: to what extent social programming of the HPA axis occur and why? To what extent these induced changes are reversible? Are these changes adaptive or do they reflect potential constraints?

## Acknowledgements

We thank all the students and field assistants in marmot catching. We are grateful to the municipality of Tignes for the use of the Santel chalet.

